# Vimentin binds to G-quadruplex repeats found at telomeres and gene promoters

**DOI:** 10.1101/2021.05.25.444966

**Authors:** Silvia Ceschi, Michele Berselli, Mery Giantin, Stefano Toppo, Barbara Spolaore, Claudia Sissi

**Affiliations:** Department of Pharmaceutical and Pharmacological Sciences, University of Padova, 35131, Padova, Italy; CRIBI Biotechnology Center (Centro di Ricerca Interdipartimentale per le Biotecnologie Innovative), University of Padova, 35131, Padova, Italy; Department of Molecular Medicine, University of Padova, 35131, Padova, Italy; Department of Comparative Biomedicine and Food Science, University of Padova, 35020, Legnaro, Italy

## Abstract

G-quadruplex (G4) structures that can form at guanine-rich genomic sites, including telomeres and gene promoters, are actively involved in genome maintenance, replication, and transcription, through finely tuned interactions with protein networks. In the present study, we identified the intermediate filament protein Vimentin as a binder with nanomolar affinity for those G-rich sequences that give rise to at least two adjacent G4 units, named G4 repeats. This interaction is supported by the N-terminal domains of soluble Vimentin tetramers. The selectivity of Vimentin for G4 repeats vs individual G4s provides an unprecedented result. Based on GO enrichment analysis performed on genes having putative G4 repeats within their core promoters, we suggest that Vimentin recruitment at these sites may contribute to the regulation of gene expression during cell development and migration, possibly by reshaping the local higher-order genome topology, as already reported for lamin B.

**Graphical abstract:** 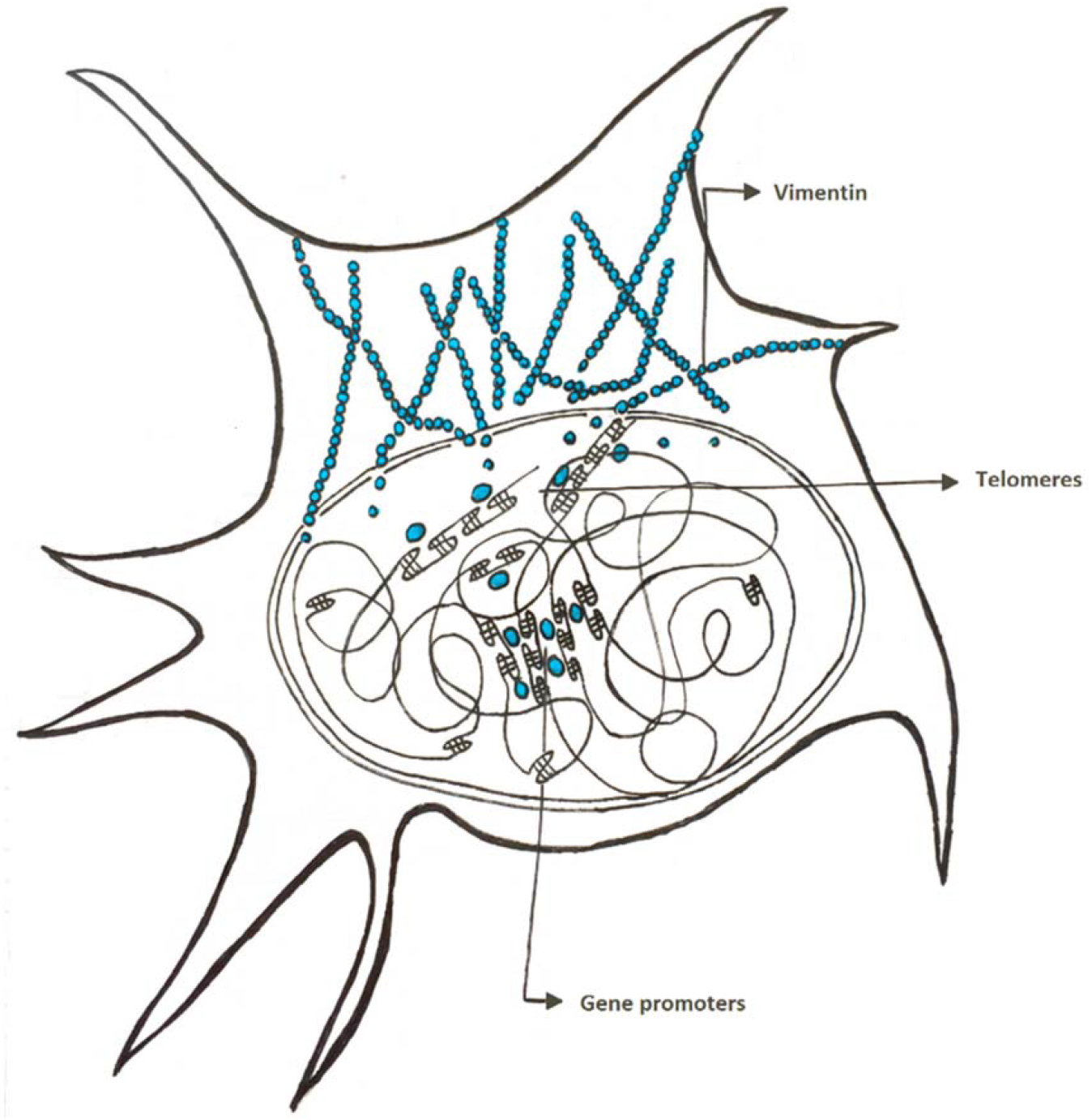

## Introduction

DNA is organized into hierarchical layers inside the nucleus. These are established through long-range chromatin interactions,^1^ which allow the creation of loops, where sets of genes are brought together into topologically associating domains (TADs)^2^ and enhancers-promoters communication is facilitated.^3^ The spatial association of TADs with similar properties, leads to the creation of the transcriptionally active (euchromatic) and repressive (heterochromatic) compartments.^4^ Heterochromatin is mainly found at the nuclear periphery, while euchromatin fills the nucleoplasm.^5^ Still, regions of active chromatin are found near nuclear pore complexes.^6^ This organization is constantly reshaped during early development and differentiation, reflecting the required gene expression program.^7^

While CCCTC binding factor (CTCF) and the cohesin complex have been identified as principally responsible for creating loops and TADs,^8^ lamin B receptor seems to play a pivotal role in tethering heterochromatin to the nuclear periphery.^9^ However, the molecular mechanisms underlying enhancers-promoters contacts and compartmentalization are still not fully elucidated. In this regard, accumulating evidence suggests that, at accessible chromatin regions,^10^ DNA folding into non-canonical structures may drive the recruitment of architectural proteins to promote gene clustering. ^11,12^

Among all DNA non-canonical secondary structures, G-quadruplexes (G4s) are tetra-helical arrangements that form within guanine-rich tracts. Here, four guanines interact through Hoogsteen hydrogen bonds to form planar arrays (G-tetrads) that stack one upon the other, building the core of the structure.^13^ G-quadruplexes can adopt different topologies depending on the relative orientation of the DNA strands (parallel, antiparallel, hybrid) and the shape of the loops connecting the G-tetrads.^14^ G-quadruplexes have been detected within the human genome.^15^ They are found at telomeres, 5’ UTR regions, introns, and gene promoters. Interestingly, G4 motifs are depleted within house-keeping genes, while they are enriched within developmental and oncogenic ones, suggesting that they may play a specific role in the regulation of these gene clusters during cell development and cancer progression.^15^ Ligands that selectively bind to G4s at promoters have been shown to influence the expression of the associated genes, thus supporting a functional role of these structures.^16^

Noteworthy, the enrichment in putative G4 forming sequences within enhancers and at TADs boundaries suggests that G-quadruplexes may regulate gene expression through their involvement in the tridimensional organization of the genome.^17^ This picture is supported by the ability of G4s to recruit architectural and chromatin remodeling factors such as the (SWI/SNF)-like chromatin remodeler ATRX,^18^ the architectural protein HMGB1,^19,20^ and the heterochromatin associated protein HP1α.^21^ Recently, Li L. and collaborators showed that G4 structures participate to the YY1-mediated DNA looping, thus providing experimental evidence to this model.^12^

Still, a clear correlation between the complex G4 structural features and protein recruitment is lacking. In the present study, we focused on peculiar G4 arrangements, to whom we will refer here as G4 repeats. G4 repeats comprise two or more adjacent G4 modules, which eventually give rise to end-to-end mutual interactions. They were first characterized for the human telomeric sequence, where multiple adjacent G4s interact through transient π-π stacking of the external tetrads.^22^ More recent studies highlighted the ability of gene promoter sequences to also give rise to G4 repeats.^23,24,25^ Among them, the hTERT sequence, located within the core promoter of telomerase, folds into three interacting parallel three-quartet G4s,^23^ the ILPR sequence, located within the promoter of insulin, folds into two cross-talking hybrid four-quartet G4s,^24^ while KIT2KIT*, which is found within the core promoter of *c-KIT*, folds into two interacting parallel three-quartet and antiparallel two-quartet G4s.^25^

With the aim to identify proteins able to interact with G4 repeats, we performed pull-down assays with KIT2KIT* using nuclear extracts from the *KIT*-positive HGC-27 cell line. Noteworthy, we found the architectural protein Vimentin as the best interactor. By using a small panel of sequences, we showed that Vimentin binds to G4 repeats regardless of their sequence and topology and with high selectivity with respect to other DNA arrangements.

Vimentin is the first reported protein selective for G4 repeats vs individual G4s. This points to G4 repeats as unique structural elements involved in the higher-order genome architecture.

## Results

### Identification of nuclear proteins that bind to the KIT2KIT* G4 repeat

To identify nuclear proteins that interact with KIT2KIT* G4 repeat, pull-down assays were performed with nuclear extracts from the *KIT*-positive HGC-27 cell line. Streptavidin-coated paramagnetic beads were derivatized with the biotinylated oligonucleotide and subsequently incubated with nuclear extracts. Bound proteins were eluted with a KCl gradient (Supplementary Fig. S1A). A last fraction was obtained by boiling beads in denaturing Laemmli sample loading buffer.^26^ When solved by SDS-PAGE (Figure 1A), this last fraction exhibited three main bands that were cut and subjected to in-gel trypsin digestion and LC-MS^E^ analyses for protein identification. Two of them (bands S1 and S2 at ^~^15 kDa and ^~^28 kDa, respectively) corresponded to Streptavidin monomer and dimer that detached from the beads along boiling procedure (Table 1; detailed data are given as Supplementary Material-Excel file). That aside, the band at ^~^50 kDa (band V in Figure 1A) was associated to the intermediate filament protein Vimentin (Table 1; detailed data are given as Supplementary Material-Excel file). Experiments performed with KIT2KIT* paired with its complementary strand revealed a different protein elution pattern (Supplementary Fig. S1B). Most importantly, no enrichment in Vimentin was observed within the last fraction (Figure 1B).

**Figure 1:**
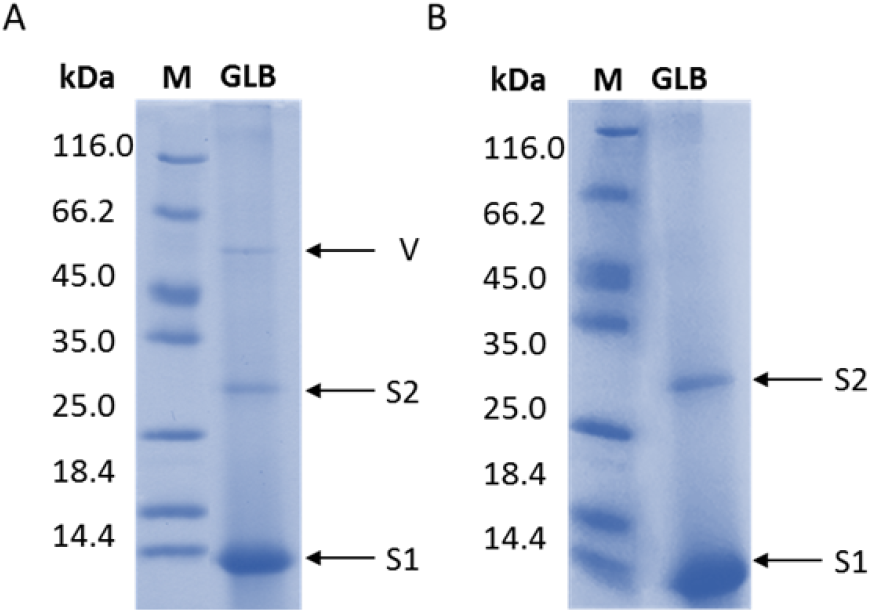
Pull-down assays identified Vimentin as a binder for the KIT2KIT* G4 repeat. SDS-PAGE of the last eluted fraction obtained from pull-down assays performed with KIT2KIT* **a** G-quadruplex and **b** duplex. Bands corresponding to Streptavidin monomer (S1) and dimer (S2) and Vimentin (V) are highlighted.

**Table 1.**
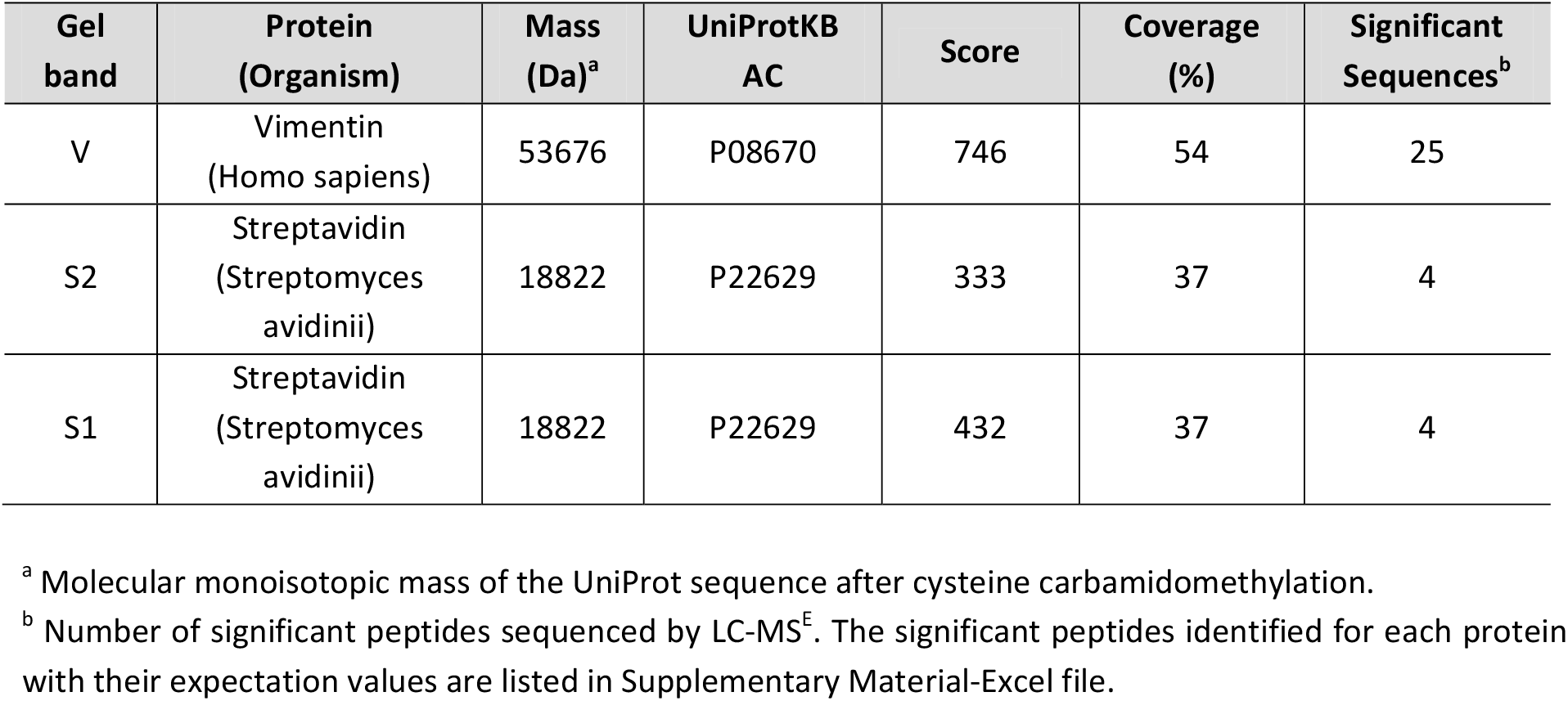
Proteins identified by LC-MS^E^.

### Vimentin selectively binds to G4 repeats

To validate the binding of Vimentin to the G4-folded KIT2KIT*, we performed electrophoretic mobility shift assays (EMSA) with the purified recombinant protein. The oligonucleotide was equilibrated in 150 mM KCl to promote G-quadruplex formation before protein addition. To avoid Vimentin polymerization into filaments, binding reactions were carried out at pH 8.4. Indeed, under these experimental conditions, Vimentin is stably arranged into tetramers.^27^ As shown in Figure 2A, at pH 8.4, free Vimentin tetramers migrate toward the anode as well as Vimentin-DNA complexes. Vimentin was proved able to bind to KIT2KIT*, leading to a well-defined band belonging to the complex. A fraction of free DNA appeared as a retarded band as a result of partial dissociation of the complex during the run. Noteworthy, no complex was observed when Vimentin was incubated with the isolated KIT2 and KIT* G-quadruplexes.

**Figure 2:**
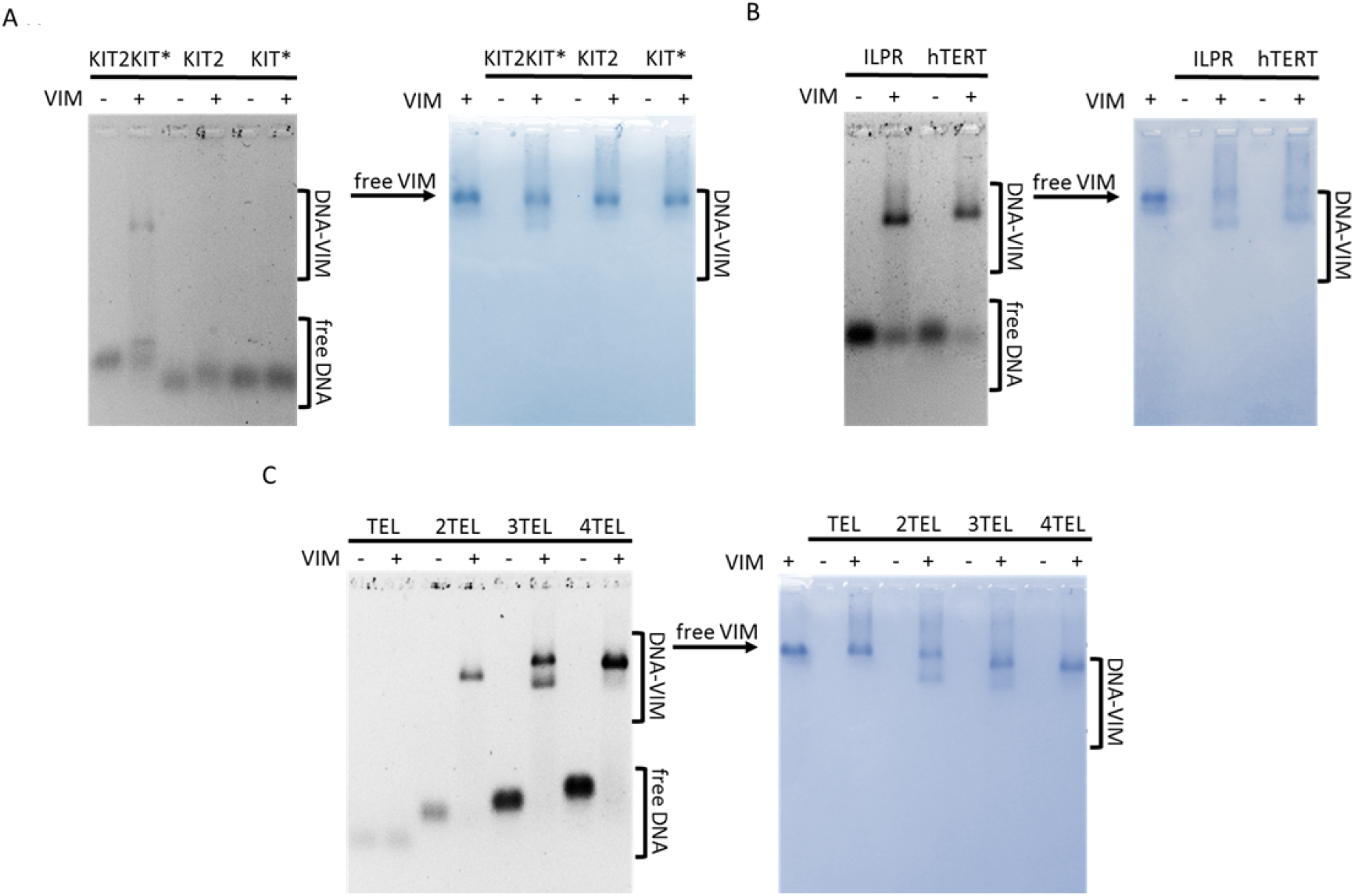
Vimentin selectively binds to G4 repeats. EMSA of 500 nM **a** KIT2KIT*, KIT2 and KIT*, **b** ILPR and hTERT, **c** TEL, 2TEL, 3TEL and 4TEL G-quadruplexes with 8 μM Vimentin in 5 mM Tris-HCl (pH 8.4), 150 mM KCl, stained with Sybr Green II (on the left) and with colloidal Coomassie Brilliant Blue G250 (on the right).

To determine whether KIT2KIT* recognition was based on sequence composition or structural features, we tested other DNA sequences for which the folding into G4 repeats akin to KIT2KIT* was already reported. Figure 2B shows EMSA performed with the insulin-linked polymorphic region (ILPR) and the human telomerase promoter (hTERT). With both oligonucleotides Vimentin formed single well-defined complexes. The heterogeneity of the so far tested sequences suggests that Vimentin recruitment is driven by DNA folding into G4 repeats, irrespectively of G-quadruplex topology and number of G-tetrads. Therefore, to better characterize this interaction, we moved to the telomeric sequence, as it constitutes an easily tunable model for G4 repeats. Indeed, by increasing the number of TTAGGG repeats, the resulting oligonucleotide folds into one (TEL), two (2TEL), three (3TEL) or four (4TEL) adjacent G-quadruplexes.^22^ Moreover, Vimentin association with telomeres has already been observed within living cells.^28^ As expected, Vimentin did not bind to the single telomeric G4 (Figure 2C) while it interacted with the G4 repeats. With 2TEL and 4TEL it formed a single complex whereas in the presence of 3TEL, two different complexes were detected, possibly reflecting the reported conformational heterogeneity of this oligonucleotide in solution.^29^

The selectivity of Vimentin for G-quadruplex vs duplex and single-stranded DNA was proved by performing EMSA with double-stranded KIT2KIT* and 2TEL and with a 49-mers G-rich oligonucleotide (G-rich noG4) unable to fold into G4, as demonstrated by circular dichroism (Supplementary Fig. S2). Vimentin little interacted with double-stranded DNA, leading to poorly defined complexes (Supplementary Fig. S3). As regards the unfolded single-stranded oligonucleotide, no binding was detected. Interestingly enough, when the same experiments were performed under KCl-free conditions, Vimentin interacted with both single and double-stranded oligonucleotides (Supplementary Fig. S3), in line with the already reported association of Vimentin with G-rich DNA.^30,31^ The fact that this interaction is completely abolished in the presence of 150 mM KCl, while the binding to G4 repeats is maintained, suggests that the binding of Vimentin to duplex and unfolded oligonucleotides is based on aspecific electrostatic interactions, while its binding to G4s relays on a more specific binding pattern.

### One Vimentin tetramer binds to two adjacent G4s with nanomolar affinity

The stoichiometry of Vimentin binding to G4 repeats was investigated according to the method of continuous variations (or Job method),^32^ the complex formed at variable Vimentin and oligonucleotide molar fractions being solved by agarose gels and quantified. The Job plot derived for the complex of Vimentin with the telomeric G4 repeat 2TEL (Figure 3A) showed maximal complex formation at 0.2 DNA molar fraction, corresponding to a 1:4 DNA:Vimentin binding stoichiometry. Thus, interaction is likely to occur between a Vimentin tetramer and two adjacent G4s. Consistently, in the presence of 4TEL, the Job plot showed a maximum at 0.1 DNA molar fraction, confirming the recruitment of two Vimentin tetramers on a stretch of four contiguous G4s (Figure 3B).

**Figure 3:**
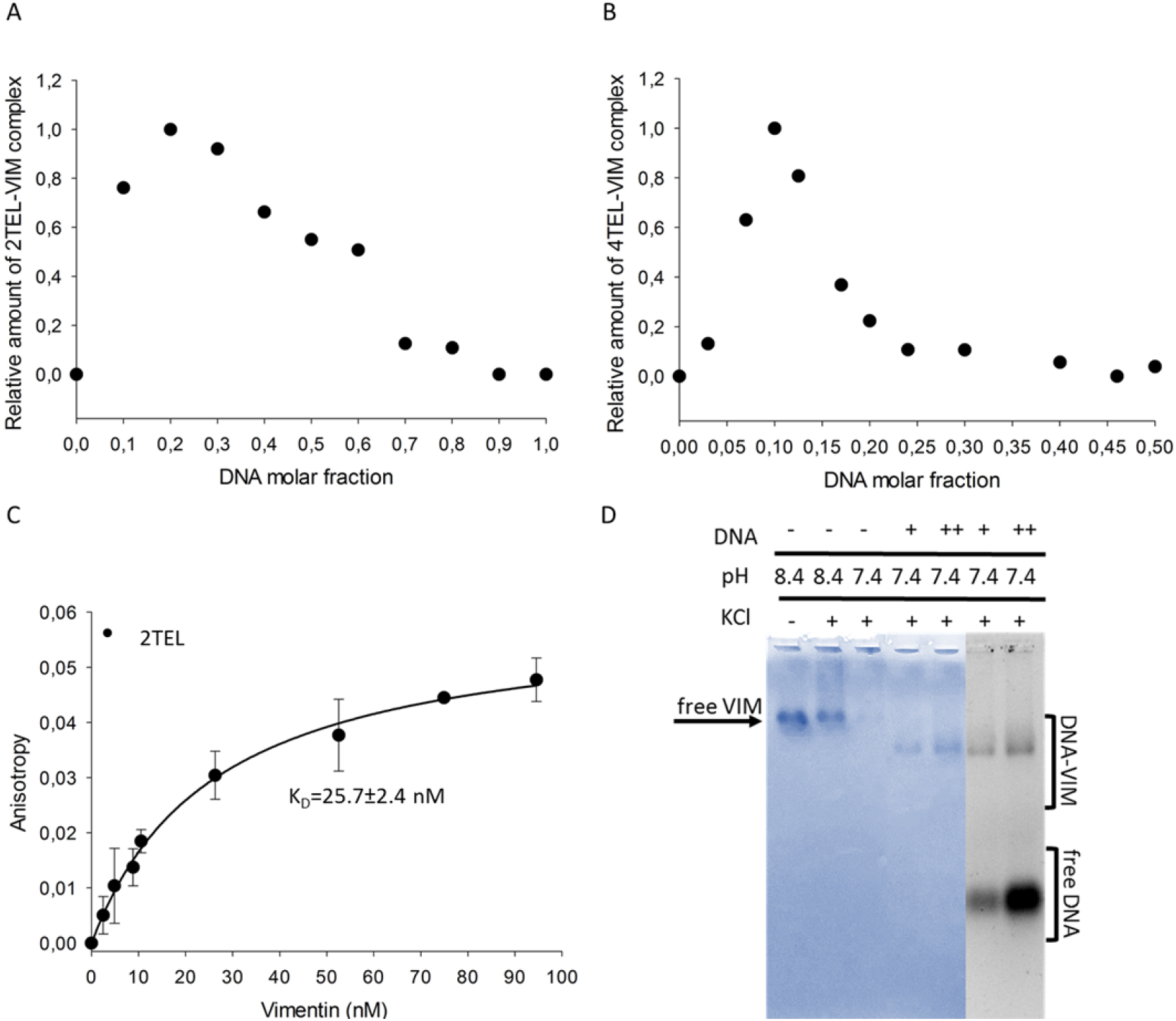
Vimentin binds to the telomeric G4 repeat as a soluble tetramer with a 1:1 stoichiometry and nanomolar affinity. Job plot derived from EMSA performed with Vimentin and **a** 2TEL, **b** 4TEL, in 5 mM Tris-HCl (pH 8.4), 150 mM KCl, at constant sum of oligonucleotide and protein concentrations (10 μM), and by varying their molar fraction. **c** Fluorescence anisotropy for monitoring the binding of Vimentin with the telomeric G4 repeat (5 nM 5’-6-FAM-2TEL) in 5 mM Tris-HCl (pH 8.4), 150 mM KCl at 25 °C. The data represent mean ± s.d. from three independent experiments. Vimentin concentration is calculated as tetramers concentration. **d** EMSA of 8 μM Vimentin with 2 μM (+) and 8 μM (++) 2TEL performed in 5 mM Tris-HCl (pH 7.4), 150 mM KCl, stained with Coomassie Brilliant Blue G250 and Sybr Green II.

To quantitatively determine the affinity of Vimentin for the telomeric G4 repeat, we followed the change in fluorescence anisotropy of 5’-6-FAM labelled 2TEL, upon titration with the protein. Analyses were performed considering the concentration of Vimentin tetramers and data were fitted according to a 1:1 binding model. Results showed that Vimentin binds strongly to 2TEL, with a *K_D_* value of 25.7 ± 2.4 nM (Figure 3C).

### Vimentin binding to G4 repeats competes with filament assembly

To assess whether Vimentin assembly into filaments impacts on its G4 binding properties, we performed EMSA with 2TEL in 150 mM KCl, at pH 7.4 (Figure 3D). Indeed, as previously reported by Herrmann and colleagues,^27^ lowering the pH to 7.4 in the presence of high ionic strength, causes extensive polymerization of Vimentin. This clearly emerges from the almost complete disappearance of the band corresponding to Vimentin tetramers in agarose gels (third lane of Figure 3D). Interestingly, when 2TEL was added to polymerized Vimentin, the complex with the soluble tetrameric form of the protein formed, as evidenced by the appearance of its characteristic band. In line with the dynamic reversible assembly of Vimentin,^33^ the protein polymerization into filaments does not prevent the binding to G4 repeats. Instead, this interaction competes with filament assembly.

### Vimentin N-terminal domain is involved in the interaction with G4 repeats

To identify the Vimentin domains that are involved in the interaction with G4 repeats, we performed limited proteolysis experiments on tetrameric Vimentin in the absence and in the presence of stoichiometric amounts of 2TEL. Proteolysis was performed with trypsin since Vimentin contains several lysine and arginine residues homogenously distributed along the sequence (Supplementary Fig. S4). Proteolysis reaction mixtures obtained after different times of incubation were solved by SDS-PAGE (Figure 4) and the sequence of the fragments was determined by in-gel trypsin digestion followed by LC-MS^E^ analyses. In Table 2 the aminoacidic sequence assigned to each band is reported.

**Figure 4:**
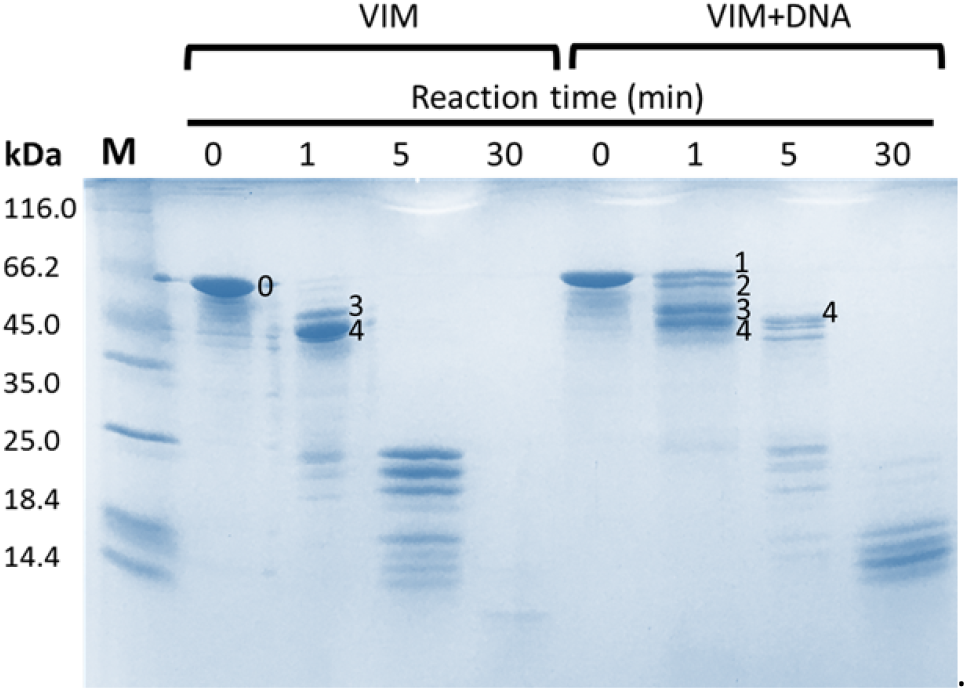
Vimentin N-terminal domain is involved in the interaction with G4 repeats. SDS-PAGE of the mixtures obtained from limited proteolysis performed with trypsin on tetrameric Vimentin, in 5 mM Tris-HCl (pH 8.4), 150 mM KCl, in the absence or in the presence of stoichiometric amounts of 2TEL. The reaction was quenched after 0, 1, 5 and 30 minutes from trypsin addition. Labelled bands were cut and subjected to ‘in-gel’ trypsin digestion and LC-MS^E^ analyses for peptide sequencing.

**Table 2.** Vimentin fragments identified in the bands by in-gel trypsin digestion and LC-MS^E^ analyses.

| Band | Fragment | Mass (kDa) |
| --- | --- | --- |
| 0 | 1–465 | 53.6 |
| 1 | 4–465 | 53.2 |
| 2 | 4–438 | 50.1 |
| 3 | 69–465 | 46.5 |
| 4 | 69–438 | 43.3 |

Analysis of the Vimentin band at time = 0 min (band 0 in Figure 4) gave a sequence coverage of 91% (Supplementary Material-Excel file), with only some internal regions missing (amino acids 97–99, 235–269, 292–293 and 310-312). In the absence of DNA, after 1 min from trypsin addition, full-length Vimentin was converted into two main species (bands 3 and 4 in Figure 4), corresponding to the protein lacking respectively the N-terminal domain (band 3), and both the C-terminal and N-terminal domains (band 4). This result fits with the intrinsically unfolded state of the N-terminal and C-terminal domains that makes them readily subjected to proteolysis. Addition of DNA promoted a delay in proteolysis. Worth of note, after 1 min from trypsin addition, two different species were generated from the full-length protein (bands 1 and 2 in Figure 4). These correspond to Vimentin lacking the first three amino acids at the N-terminal domain (band 1) and to Vimentin lacking the same three amino acids at the N-terminal domain plus the C-terminal domain (band 2). These data confirm the involvement of the N-terminal domain in the binding to G4 repeats.

### Putative G4 repeats are found within the promoter of genes involved in the cellular response to external stimuli, cell-cell communication and locomotion

So far, we showed that Vimentin selectively binds to G4 repeats. To search for sequences putatively able to adopt such conformation, we previously developed QPARSE tool and highlighted their non-random distribution within human gene promoters.^34^ Here, we refined our search focusing on the first 100bp upstream the Transcription Starting Site (TSS) of the genes, since this is the region where we previously found the highest frequency in putative G4 repeats.^34^ Moreover, it comprises the already characterized G4 repeats KIT2KIT* and hTERT, for which a role in controlling the expression of the downstream gene has been experimentally confirmed.^35,36^

All the promoter regions corresponding to genes annotated in GENCODE v34 (38404 sequences) were downloaded from ENSEMBL.^37^ The software identified 1477 genes containing at least one putative double G4 repeat (two adjacent G4s) and 295 sequences containing a putative triple G4 repeat (three adjacent G4s). Sequences with a triple G4 repeat were a subset of those with a double G4 repeat, as expected, apart for only one gene due to the slightly different searching criteria. Overall, the retrieved sequences potentially able to fold into a double or triple G4 repeat are 1478. The median GC content of these sequences is 80% and 98% of this subset shares a GC content greater than 60%. We refer to this list as double_triple_G4_PQS. We further selected a background population for comparison including 14053 sequences that share the same high GC content with double_triple_G4_PQS (greater than 60%) but do not contain any putative double or triple G4 repeat. We call this list GC_rich_BKG.

Using these two lists of genes, we performed a GO enrichment analysis using DAVID tool,^38^ to look for a link between the reported physio-pathological roles of the G4 repeats containing genes, and those related to Vimentin.

The more interesting results are summarized in Figure 5 (Detailed results are found in Supplementary Material-Excel file).

**Figure 5:**
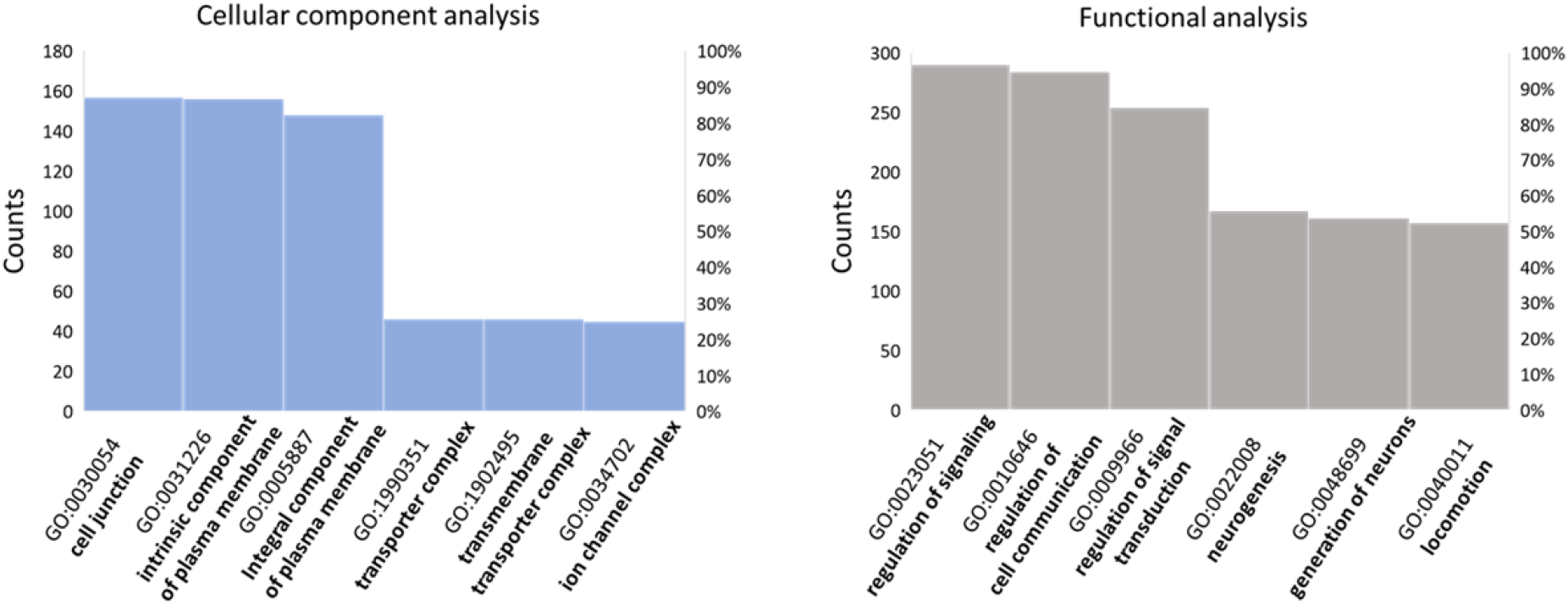
Putative G4 repeats are enriched within the core promoters of genes involved in the cellular response to external stimuli, cell-cell communication, and locomotion. Results of GO enrichment analysis performed with David tool on human genes presenting putative G4 repeats within their promoters.

Cellular component analysis revealed a significant enrichment in cell membrane components, particularly in those engaged in cell junctions. Both biological process and functional analyses highlighted an enrichment in proteins responsible for cell-cell communication, signal transduction and locomotion, together with an overrepresentation of genes involved in neurogenesis and nervous system development. We performed the same analyses with Panther tool^39^ and again we found a significant enrichment in plasma membrane components, particularly those participating to cell surface signaling, cellular response to external stimuli and cell-cell communication.

## Discussion

Vimentin is an intermediate filament protein highly expressed within migratory cells that are present at the early stage of embryonic development.^40^ Its postnatal expression is restricted to motile cells such as fibroblasts, endothelial cells, lymphocytes, and Swann cells.^41^ Noteworthy, epithelial cells rely on Vimentin expression to acquire fibroblast-like morphology and increased migratory capacity during epithelial to mesenchymal transition (EMT), which occurs both during physiological tissue development/regeneration and pathological cancer progression toward metastasis.^42^

As a structural protein with main cytosolic localization, Vimentin function reported so far is to orchestrate cytoskeletal rearrangements and mechano-signaling in support to cell migration.^41^ Interestingly, studies conducted on poorly differentiated metastatic cancer cells revealed the presence of Vimentin within their nuclear matrixes, while it was no longer detected upon induction of cell differentiation.^43,44^ Moreover, in human embryo fibroblasts, Vimentin has been found in tight association with telomeres and centromeres.^28^ These data point to specific functions of Vimentin at nuclear level.

In the present study, we found that the fraction of Vimentin that was present within nuclear extracts of undifferentiated HGC-27 cells, binds to DNA in a structure dependent/sequence independent manner. Indeed, it efficiently interacts with different G4-folded DNA sequences with almost no binding to the corresponding unfolded/duplex conformations. As a further level of specificity, the presence of at least two adjacent G4s is required for Vimentin recruitment, regardless of the topology of the participating G4s (parallel+antiparallel for KIT2KIT*, hybrid for telomeric G4s and ILPR, parallel for hTERT) and the total number of G-tetrads (five for KIT2KIT*, six for 2TEL, eight for ILPR, nine for hTERT). Indeed, Vimentin does not bind to individual G4s, regardless of their number of G-tetrads (two for KIT*, three for KIT2 and TEL) and topology (antiparallel for KIT*, parallel for KIT2 and hybrid for TEL). This is the first time to our knowledge that a G4-binding protein displays selectivity for G4 repeats.

Vimentin exists in a highly dynamic state within living cells, where post-translational modifications drive filament assembly/disassembly, in response to changes that occur in the extracellular environment.^33,45^ In the present study, we showed that Vimentin binds to DNA G4 repeats in the tetrameric form, the stoichiometry of the complexes being one Vimentin tetramer every two adjacent G4s. Noteworthy, this interaction shifts the Vimentin assembly equilibrium toward the naturally present soluble fraction.^46^ Vimentin assembly into filaments proceeds through the lateral association of Vimentin tetramers into unit length filaments (ULF) followed by the N-terminal to C-terminal longitudinal annealing of ULF to yield mature filaments.^47^ Our limited proteolysis experiments showed that the interaction of Vimentin with G4s occurs at the N-terminal domains, and this can affect the longitudinal annealing of ULF into filaments. Noteworthy, Vimentin tetramers switch from A11-type (where the N-terminal domains are oriented toward the center of the tetramer) to A22-type (where the N-terminal domains are placed at the edge of the tetramer) during ULF formation.^48^ Therefore, the binding of G4 repeats to the soluble A11-type tetramers may also prevent the type-switching and, consequently, ULF formation.

The high affinity of Vimentin for G4 repeats fits with its already reported association with telomeres and centromeres. The herein acquired *in vitro* evidence of Vimentin binding to G4 repeats at gene promoters suggests that the same interaction may occur within living cells as well. In this regard, GO enrichment analysis performed on genes having putative G4 repeats within their core promoters revealed an overrepresentation of cell membrane components, particularly those participating to cell-cell communication and cell surface signaling. Noteworthy, among them there is the zinc finger protein SNAI1, the expression of which was shown to be directly regulated by Vimentin during EMT.^42^ It is thus tempting to suggest that the binding of Vimentin at G4 repeats may contribute to the regulation of the expression of the associated genes, possibly contributing to wider DNA topological changes. In this regard, Vimentin was shown to influence not only nuclear shape and mechanics, but also chromatin condensation within mesenchymal cells.^49^ It has been reported that soluble pools of Vimentin-related lamin A/C and B can contact euchromatin within the nucleoplasm, promoting gene clustering within the so-called euchromatin lamin associated domains (eLADs), ultimately regulating their expression.^50,51^ Of particular interest is the involvement of lamin B1-eLADs in chromatin reorganization that occurs during EMT. Indeed, lamin B1 was shown to bind to G-rich promoters of genes that belong to the EMT pathway, helping the establishment of the EMT transcriptional program.^51^ Soluble pools of Vimentin may exert a similar function, or even provide lamin B a way to contact DNA at G-rich promoters. Indeed, the direct interaction of Vimentin C-terminal domain with lamin B has already been reported *in vitro*.^52^

GO enrichment analysis also highlighted the presence of putative G4 repeats within the promoters of genes involved in neurogenesis and nervous system development. In this regard, the relevance of Vimentin in neurological development is well established.^53,54^ In a recent study, high levels of soluble Vimentin were detected in the axoplasm of neurons following neuronal injury. Vimentin was found able to translocate from the site of injury to the soma through direct interaction with β-importin and dynein-mediated retrograde transport. The authors point to a signaling role for soluble Vimentin during neural injury, in support to neurite regeneration.^55^ Noteworthy, the direct interaction of Vimentin with β-importin, provides a mechanism for its entry into the nucleus in response to peripheral stimuli.

Overall, these evidences support a correlation between the functions of soluble Vimentin and those of the genes containing putative Vimentin binding sites at their promoters.

To conclude, in the present study, we identified the intermediate filament protein Vimentin as a selective binder for G4 repeats. The fact that Vimentin does not bind to individual isolated G4s could provide her a way to contact DNA at specific genomic loci, including telomeres, centromeres, and a distinct subset of gene promoters. Further studies are needed to unravel the biological significance of such interaction within human living cells. Our working hypothesis is that G4 repeats may exist as primary structural elements, able to drive the recruitment of architectural proteins to ultimately reshape the higher-order genome folding during important physiological processes such as cell development, differentiation and migration, thus favoring the establishment of the required gene expression program.

## Methods

### Oligonucleotides

Oligonucleotides were purchased lyophilized and RP-HPLC purified from Eurogentec (Seraing, Belgium) and used without further purification. The DNA sequences are listed in Table 3.

**Table 3.**
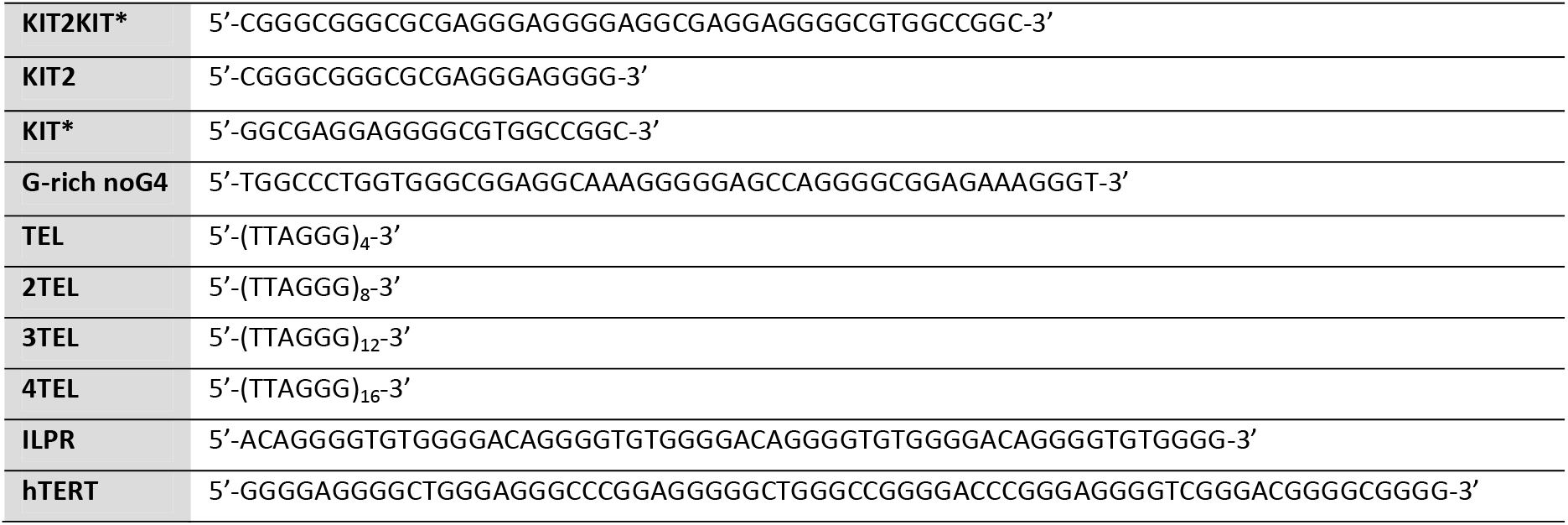
DNA sequences.

Oligonucleotides were resuspended in nuclease-free water from Thermo Fisher Scientific (Waltham, MA, USA) to obtain 100 μM stock solutions, which were then diluted in the proper buffer for further analyses. The concentrations of the initial stock solutions were measured by UV absorbance at 260 nm on a Uvikon XS, using molar absorption coefficients calculated with a nearest neighbour model.^56^

The solutions were annealed with 85 °C heating for 7 min and then led to equilibrate overnight at room temperature. Equilibrated solutions were doped with KCl to promote G-quadruplex folding. For duplex DNA preparation, oligonucleotides were annealed in the presence of equimolar amounts of the complementary strand.

### Protein sample preparation

Recombinant human Vimentin was purchased lyophilized and RP-HPLC purified from LS-Bio (Seattle, WA, USA) and was stored at −80°C in 8 M urea, 5 mM Tris-HCl (pH 7.5), 1 mM dithiothreitol, 1 mM EDTA, 0.1 mM EGTA. The day before use, Vimentin was renatured following a protocol developed by Herrmann and colleagues to avoid extensive polymerization into filaments.^27^ Briefly, the protein was dialyzed at room temperature against dialysis buffer (5 mM Tris-HCl (pH 8.4), 1 mM EDTA, 0.1 mM EGTA, and 1 mM dithiothreitol) containing progressively reduced urea concentration (6, 4, 2, and 1 M urea). Dialysis against a large volume of dialysis buffer was continued overnight at 4°C. The next day, dialysis was continued into tetramer buffer (5 mM Tris-HCl (pH 8.4)) for 1 h at room temperature. After dialysis, the concentration of Vimentin monomers was determined by measuring the absorption at 280 nm with ε = 24,900 cm ^-1^M^-1^.

### Nuclear extracts

The human gastric carcinoma cell line HGC-27 (European Collection of Authenticated Cell Cultures (ECACC) 94042256) was grown in Eagle’s Minimal Essential Medium (EMEM, Gibco^®^ Life Technologies, Carlsbad, USA) supplemented with 10% fetal bovine serum (Gibco^®^ Life Technologies), 2 mM L-glutamine (Euroclone, Milan, Italy), 1% non-essential amino acids (Euroclone) and 1% penicillin/streptomycin (Euroclone), under a humidified 5% CO_2_ atmosphere, at 37°C. Nuclear extracts were obtained as described by Abmayr and Workman with minor changes.^57^ Briefly, confluent HGC-27 cells, previously seeded in 4 Petri dishes (10-cm diameter) were scraped off and incubated with a hypotonic buffer (20 mM HEPES (pH 7.9), 1.5 mM MgCl_2_, 20 mM KCl, 0.5 % Triton X-100, 0.5 mM PMSF, 1 mM dithiothreitol, 1X protease inhibitor cocktail (Merck, Darmstadt, Germany)) for 15 min on ice. Nuclei were pelleted by centrifugation and then incubated with a high-salt buffer (20 mM HEPES (pH 7.9), 25% glycerol, 1.5 mM MgCl_2_, 420 mM KCl, 0.5 % Triton X-100, 0.5 mM PMSF, 1 mM dithiothreitol, 1X protease inhibitor cocktail) for 30 min on ice. Released nuclear proteins were quantified according to the Bradford method.

### Pull-down Assays

600 μL of Streptavidin-coated paramagnetic particles (Promega, Milan, Italy) were washed three times with 600 μL of PBS-1X (Euroclone, Milan, Italy) and then resuspended in 60 μL of PBS-1X. 2 μM 5’-biotinylated KIT2KIT* was previously annealed in 10 mM potassium phosphate (pH 7.4) and subsequently equilibrated overnight in the presence of 150 mM KCl, to promote G-quadruplex folding. For duplex DNA preparation, 2 μM 5’-biotinylated KIT2KIT* was annealed in 10 mM potassium phosphate (pH 7.4) in the presence of equimolar amounts of the complementary strand.

The oligonucleotide was then added to the beads and incubation was performed for 30 min at room temperature. Beads were washed three times with 600 μL of oligonucleotide buffer and then resuspended in 60 μL of pull-down buffer (20 mM HEPES (pH 7.9), 100 mM KCl, 0.2 mM EDTA, 0.5 mM dithiothreitol, 0.2 mM PMSF, 3 μM poly[dA-dT], 20% glycerol). 100 μg of nuclear extract were added to the beads and incubation was performed for 1 h at room temperature. Protein elution was performed with 60 μL of pull-down buffer containing increasing concentration of KCl (250 mM, 500 mM, 750 mM, 1M). The last elution was performed by boiling beads for 5 min in Laemmli SDS-PAGE sample loading buffer.^26^ All fractions were run on Laemmli SDS-PAGE. After staining with colloidal Coomassie Brilliant Blue G250, interesting bands were cut and subjected to in-gel trypsin digestion and LC-MS^E^ analyses for protein identification.

### Electrophoretic Mobility Shift Assays

Electrophoretic mobility shift assays (EMSA) were performed on 1.5 % agarose gels in 0.5X TBE buffer (0.89 M Tris-Borate, 20 mM EDTA, pH 8.0) supplemented with 10 mM KCl. Samples contained 500 nM DNA and 8 μM Vimentin, in 5 mM Tris-HCl (pH 8.4), 150 mM KCl. For the Job method, samples contained constant sum of Vimentin and oligonucleotide concentrations (10 μM), at variable molar fractions. Vimentin was added on oligonucleotides previously annealed in the reaction buffer and incubation was performed for 1 h at room temperature. Gel loading buffer (50% glycerol, 50% water) was added to the samples immediately before loading. Electrophoresis proceeded for 1 h at 6V/cm at 4 °C. Gels were stained with Sybr Green II for nucleic acid visualization and, afterwards, with Coomassie Brilliant Blue G250 for protein detection. Gel images were acquired with a Geliance 600 apparatus. Bands belonging to the DNA-Vimentin complex were quantified with ImageJ software. Quantification performed on Sybr Green II or Coomassie stained gels provided identical results.

### Circular Dichroism Spectroscopy

Circular dichroism (CD) spectra were acquired on a JASCO J-810 spectropolarimeter equipped with a Peltier temperature controller. CD spectra were recorded from 235 to 330 nm with the following parameters: scanning speed of 100 nm/min, band width of 2 nm, data interval of 0.5 nm and response of 2 s. Measurements were performed using a 1 cm path length quartz cuvette at oligonucleotide concentration of 2 μM in 5 mM Tris-HCl (pH 8.4), 150 mM KCl. Observed ellipticities were converted to Molar Ellipticity which is equal to deg·cm^2^·dmol^-1^, calculated using the DNA residue concentration in solution.

### Fluorescence anisotropy

Fluorescence anisotropy measurements were performed on a JASCO FP-6500 spectrofluorometer equipped with polarization devices and with a Peltier temperature controller. Measurements were performed at 25°C using a 1 cm path length quartz cuvette with the following parameters: 495 nm excitation wavelength, 520 nm emission wavelength, band width of 5 nm, response 8s, sensitivity high, 2 acquisitions. The instrument G factor was determined prior to anisotropy measurements. 5’-6-FAM labelled oligonucleotides were used at a concentration of 5 nM in 5 mM Tris-HCl (pH 8.4), 150 mM KCl. Titrations were performed by adding increasing concentrations of recombinant Vimentin to the oligonucleotide solution. After mixing, the solution was led to equilibrate for 10 min at room temperature before acquisition. Experiments were performed in triplicate. Acquired data were fitted according to a 1:1 binding model with the following equation:

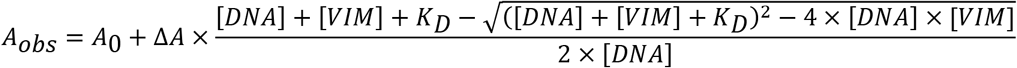

where [DNA] and [VIM] stand for DNA and tetrameric Vimentin concentrations, respectively; *A*_obs_ is the observed anisotropy value; *A*_0_ is the anisotropy value in the absence of protein; Δ*A* represents the total change in anisotropy between free and fully bound DNA, and *K*_D_ is the equilibrium dissociation constant.

### Trypsin limited proteolysis

The solution of Vimentin in 5 mM Tris-HCl (pH 8.4) was loaded on a Pierce™ Detergent Removal Spin Column (Thermo Scientific) and eluted in the same buffer. Limited proteolysis was conducted on this protein solution (0.43 mg/mL) without or with addition of stoichiometric amounts of DNA, in 150 mM KCl. The reaction was initiated upon addition of trypsin at an enzyme/substrate ratio of 1/500 by weight and incubated at 37 °C without stirring for 1, 5 and 30 min. Aliquots taken at each time of incubation were quenched by adding 0.3% TFA solution, lyophilized and then analyzed by SDS-PAGE. Labelled bands were cut and subjected to ‘in-gel’ trypsin digestion and LC-MS^E^ analyses for peptide sequencing. The sequence of full-length Vimentin used in the experiments was also confirmed by MS analysis of the recombinant protein (calculated average mass 53520.6 Da and measured mass 53520.6 Da, Supplementary Fig. S5).

### In-gel digestion

In-gel digestion of protein bands was performed according to Shevchenko et al.^58^ Briefly, excised bands were cut into small cubes, washed with water, with 50% acetonitrile in water and shrunk with neat acetonitrile. Gel particles were swelled in 10 mM dithiothreitol, 0.1 M NH_4_HCO_3_ and incubated for 45 min at 56°C. After cooling at room temperature, the supernatant was replaced with the same volume of iodoacetamide solution (55 mM iodoacetamide in 0.1 M NH_4_HCO_3_) and the tubes were incubated for 30 min in the dark at room temperature. After removal of the iodoacetamide solution, gel pieces were washed again with water followed by 50% acetonitrile in water and shrunk with neat acetonitrile to remove completely the Coomassie staining. The gel particles were eventually rehydrated on ice in a solution containing 5 ng/μL of trypsin (Promega, modified sequencing grade) in 50 mM NH_4_HCO_3_. After complete rehydration, gel pieces were covered with 50 mM NH_4_HCO_3_ and incubated overnight at 37°C. The supernatants were then transferred to clean tubes and peptides were extracted from gel particles upon incubation with 5 % formic acid in water followed by dilution with an equal volume of neat acetonitrile. All the peptide-containing supernatants were combined and dried using a Speed-Vac system (Savant).

### Mass spectrometry analyses

The tryptic digests of the gel bands were analysed using a Xevo G2-S QTof (Waters) equipped with a Waters Acquity H-Class UPLC system. Mobile phase A consisted in 0.1% formic acid in water while mobile phase B was 0.1% formic acid in acetonitrile. For protein identification, LC analyses were performed using an ACQUITY BEH C18 VanGuard Pre-Column (2.1×5mm, 1.7μm, Waters) in line with an ACQUITY UPLC BEH C18 column (1.0×100mm, 1.7μm, Waters). Peptide separation was performed using a linear gradient from 3% to 65% of B in 36 minutes at a flow rate of 0.1 mL/min, with a column temperature set at 30 °C. For the digests of the bands from the proteolysis experiments, separations were carried out on an AdvanceBio Peptide Map Guard (2.1×5 mm, 2.7μm, Agilent technologies) and AdvanceBio Peptide Map column (2.1×150 mm, 2.7μm). Peptide separation was performed using a linear gradient from 2% to 65% of B in 36 minutes at a flow rate of 0.2 mL/min, with a column temperature set at 30 °C. For all the LC-MS^E^ analyses, the Xevo G2-S QTof operated in the ESI positive ion resolution mode and with a detection window between 50-2000 m/z. MS^E^ acquisition was performed by alternating two MS data functions: one for acquisition of peptide mass spectra with the collision cell at low energy (6 V), and the second for the collection of peptide fragmentation spectra with the collision cell at elevated energy (linear ramp 20 to 40 V). Analyses were performed with LockSpray™ using a solution of 1 ng/μL LeuEnk in 50% acetonitrile, 0.1% formic acid in water. For protein identification, LC-MS^E^ data were analysed using the on-line MASCOT server (www.matrixscience.com) against the Swiss-Prot database (release 2020_06). For Vimentin identification, a taxonomy filter to Homo sapiens was applied and a contaminants database was included (See Supplementary Material-Excel file).

The following parameters were used in the MASCOT search: trypsin specificity; maximum number of missed cleavages, 1; fixed modification, carbamidomethyl (Cys); variable modifications, oxidation (Met); peptide mass tolerance, ± 10 ppm; fragment mass tolerance, ± 15 ppm; protein mass, unrestricted; mass values, monoisotopic. A protein was considered identified when two unique peptides with statistically significant scores (p < 0.05) were obtained.

MS^E^ data of the gel bands from the proteolysis experiment were processed with the BiopharmaLynx, setting trypsin as digest reagent, 1 missed cleavage and carbamidomethyl cysteine as fixed modification. MS ion intensity threshold was set to 100 counts, and the MS^E^ threshold was set to 100 counts. MS mass match tolerance and MS^E^ mass match tolerance were set to 10 ppm. The peptide list obtained from the LC-MS analysis of Vimentin at 0 min of proteolysis was reduced to represent only peptides with an intensity higher than 3000 counts and it was considered as the reference list of tryptic peptides. For the digests of the bands at the different times of incubation, peptides with a signal higher than 100 counts were considered identified, provided that they displayed a retention time in accordance with the same peptide in the reference list. In order to identify the region of Vimentin in the different gel bands, the percent ratio between the intensity of each peptide in the LC-MS analysis of the digest of the band and in the reference list was calculated. This calculation was performed in order to consider the different ionization efficiency of the different peptides.

### QPARSE search and GO enrichment analysis

The whole set of human genes were downloaded from ENSEMBL^37^ (GENCODE v34) and the upstream 100 nucleotides from the transcription starting site (TSS) of each gene were extracted to search for double and triple putative G4 repeats (double_triple_G4_PQS). The pattern search was performed using QPARSE^34^ with the following options: i) for double G4 repeats we searched for islands of at least 3 (-m 3) and up to 4 (-M 4) Gs/Cs with connecting loops of maximum 5 nucleotides (-L 5), and at least 5 perfect islands (-p 5) out of 8 islands (-n 8) with the bulged islands that contain only one gap of length 1 (-l 1), ii) for triple G4 repeats we searched for islands of at least 3 (-m 3) and up to 4 (-M 4) Gs/Cs with connecting loops of maximum 5 nucleotides (-L 5), and at least 8 perfect islands (-p 8) out of 12 islands (-n 12) with the bulged islands that contain only one gap of length 1 (-l 1). The searched pattern was extended both in the forward and reverse strand (using the parameter -b C and -b G to search for PQS in the reverse and forward strand respectively). For comparison purposes, we calculated the GC content of the 100bp upstream the TSS of all the sequences. To assess whether genes containing putative G4 repeats were enriched in functional categories or signaling pathways, we selected genes with a high GC content similar to that found in genes containing the searched motifs. We used this list as background population. The list of double_triple-G4_PQS vs. the background population were analyzed in DAVID tool.^38^

## Acknowledgements

This research was funded by University of Padova, grant number # SISS_SID19_01. S.C. PhD fellowship was founded by University of Padova.

## Author contributions

S.C, S.T. and C.S. conceived the project; S.C. acquired the experimental data; B.S. performed proteolysis reactions and mass spectrometry analyses; S.C. and C.S analyzed experimental data. M.B. and S.T. performed bioinformatic analyses; M.G. supervised cell cultures and nuclear extracts production; C.S. raised the funds. S.C and C.S. wrote the manuscript, which was reviewed and commented by all co-authors.

## References

1. Lieberman-Aiden, E. et al. Comprehensive mapping of long-range interactions reveals folding principles of the human genome. Science 326, 289–293 (2009).

2. Dixon, J. R. et al. Topological domains in mammalian genomes identified by analysis of chromatin interactions. Nature 485, 376–380 (2012).

3. Symmons, O. et al. The Shh Topological Domain Facilitates the Action of Remote Enhancers by Reducing the Effects of Genomic Distances. Dev. Cell 39, 529–543 (2016).

4. Schoenfelder, S. & Fraser, P. Long-range enhancer-promoter contacts in gene expression control. Nat. Rev. Genet. 20, 437–455 (2019).

5. Solovei, I., Thanisch, K. & Feodorova, Y. How to rule the nucleus: divide et impera. Curr. Opin. Cell Biol. 40, 47–59 (2016).

6. Buchwalter, A., Kaneshiro, J. M. & Hetzer, M. W. Coaching from the sidelines: the nuclear periphery in genome regulation. Nat. Rev. Genet. 20, 39–50 (2019).

7. Dixon, J. R. et al. Chromatin architecture reorganization during stem cell differentiation. Nature 518, 331–336 (2015).

8. Mirny, L. A., Imakaev, M. & Abdennur, N. Two major mechanisms of chromosome organization. Curr. Opin. Cell Biol. 58, 142–152 (2019).

9. Falk, M. et al. Heterochromatin drives compartmentalization of inverted and conventional nuclei. Nature 570, 395–399 (2019).

10. Shen, J. et al. Promoter G-quadruplex folding precedes transcription and is controlled by chromatin. Genome Biol. 22, 143 (2021).

11. Achar, Y. J., Adhil, M., Choudhary, R., Gilbert, N. & Foiani, M. Negative supercoil at gene boundaries modulates gene topology. Nature 577, 701–705 (2020).

12. Li, L. et al. YY1 interacts with guanine quadruplexes to regulate DNA looping and gene expression. Nat. Chem. Biol. 17, 161–168 (2021).

13. Sen, D. & Gilbert, W. Formation of parallel four-stranded complexes by guanine-rich motifs in DNA and its implications for meiosis. Nature 334, 364–366 (1988).

14. Spiegel, J., Adhikari, S. & Balasubramanian, S. The Structure and Function of DNA G-Quadruplexes. Trends Chem. 2, 123–136 (2020).

15. Hänsel-Hertsch, R. et al. G-quadruplex structures mark human regulatory chromatin. Nat. Genet. 48, 1267–1272 (2016).

16. Kosiol, N., Juranek, S., Brossart, P., Heine, A. & Paeschke, K. G-quadruplexes: a promising target for cancer therapy. Mol. Cancer 20, 40 (2021).

17. Hou, Y. et al. Integrative characterization of G-Quadruplexes in the three-dimensional chromatin structure. Epigenetics 1–18 (2019) doi:10.1080/15592294.2019.1621140.

18. Law, M. J. et al. ATR-X Syndrome Protein Targets Tandem Repeats and Influences Allele-Specific Expression in a Size-Dependent Manner. Cell 143, 367–378 (2010).

19. Amato, J. et al. HMGB1 binds to the KRAS promoter G-quadruplex: a new player in oncogene transcriptional regulation? Chem. Commun. Camb. Engl. 54, 9442–9445 (2018).

20. Amato, J. et al. Insights into telomeric G-quadruplex DNA recognition by HMGB1 protein. Nucleic Acids Res. 47, 9950–9966 (2019).

21. Roach, R. J. et al. Heterochromatin protein 1α interacts with parallel RNA and DNA G-quadruplexes. Nucleic Acids Res. 48, 682–693 (2020).

22. Monsen, R. C., Chakravarthy, S., Dean, W. L., Chaires, J. B. & Trent, J. O. The solution structures of higher-order human telomere G-quadruplex multimers. Nucleic Acids Res. (2021) doi:10.1093/nar/gkaa1285.

23. Monsen, R. C. et al. The hTERT core promoter forms three parallel G-quadruplexes. Nucleic Acids Res. 48, 5720–5734 (2020).

24. Schonhoft, J. D. et al. Direct experimental evidence for quadruplex-quadruplex interaction within the human ILPR. Nucleic Acids Res. 37, 3310–3320 (2009).

25. Rigo, R. & Sissi, C. Characterization of G4-G4 Crosstalk in the c-KIT Promoter Region. Biochemistry 56, 4309–4312 (2017).

26. Laemmli, U. K. Cleavage of Structural Proteins during the Assembly of the Head of Bacteriophage T4. Nature 227, 680–685 (1970).

27. Mücke, N. et al. Assembly Kinetics of Vimentin Tetramers to Unit-Length Filaments: A Stopped-Flow Study. Biophys. J. 114, 2408–2418 (2018).

28. Tolstonog, G. V., Mothes, E., Shoeman, R. L. & Traub, P. Isolation of SDS-stable complexes of the intermediate filament protein vimentin with repetitive, mobile, nuclear matrix attachment region, and mitochondrial DNA sequence elements from cultured mouse and human fibroblasts. DNA Cell Biol. 20, 531–554 (2001).

29. Petraccone, L. et al. Structure and Stability of Higher-Order Human Telomeric Quadruplexes. J. Am. Chem. Soc. 133, 20951–20961 (2011).

30. Shoeman, R. L. & Traub, P. The in vitro DNA-binding properties of purified nuclear lamin proteins and vimentin. J. Biol. Chem. 265, 9055–9061 (1990).

31. Tolstonog, G. V., Li, G., Shoeman, R. L. & Traub, P. Interaction in vitro of type III intermediate filament proteins with higher order structures of single-stranded DNA, particularly with G-quadruplex DNA. DNA Cell Biol. 24, 85–110 (2005).

32. Job, P. Formation and stability of inorganic complexes in solution. Ann Chim 9, (1928).

33. Snider, N. T. & Omary, M. B. Assays for Posttranslational Modifications of Intermediate Filament Proteins. Methods Enzymol. 568, 113–138 (2016).

34. Berselli, M., Lavezzo, E. & Toppo, S. QPARSE: searching for long-looped or multimeric G-quadruplexes potentially distinctive and druggable. Bioinforma. Oxf. Engl. 36, 393–399 (2020).

35. Da Ros, S. et al. G-Quadruplex Modulation of SP1 Functional Binding Sites at the KIT Proximal Promoter. Int. J. Mol. Sci. 22, (2020).

36. Palumbo, S. L., Ebbinghaus, S. W. & Hurley, L. H. Formation of a unique end-to-end stacked pair of G-quadruplexes in the hTERT core promoter with implications for inhibition of telomerase by G-quadruplex-interactive ligands. J. Am. Chem. Soc. 131, 10878–10891 (2009).

37. Howe, K. L. et al. Ensembl 2021. Nucleic Acids Res. 49, D884–D891 (2021).

38. Huang, D. W., Sherman, B. T. & Lempicki, R. A. Systematic and integrative analysis of large gene lists using DAVID bioinformatics resources. Nat. Protoc. 4, 44–57 (2009).

39. Mi, H. et al. PANTHER version 16: a revised family classification, tree-based classification tool, enhancer regions and extensive API. Nucleic Acids Res. 49, D394–D403 (2021).

40. Franke, W. W., Grund, C., Kuhn, C., Jackson, B. W. & Illmensee, K. Formation of cytoskeletal elements during mouse embryogenesis. III. Primary mesenchymal cells and the first appearance of vimentin filaments. Differ. Res. Biol. Divers. 23, 43–59 (1982).

41. Battaglia, R. A., Delic, S., Herrmann, H. & Snider, N. T. Vimentin on the move: new developments in cell migration. F1000Research 7, 1796 (2018).

42. Strouhalova, K. et al. Vimentin Intermediate Filaments as Potential Target for Cancer Treatment. Cancers 12, (2020).

43. Spencer, V. A., Samuel, S. K. & Davie, J. R. Nuclear matrix proteins associated with DNA in situ in hormone-dependent and hormone-independent human breast cancer cell lines. Cancer Res. 60, 288–292 (2000).

44. Zhao, C.-H. & Li, Q.-F. Altered profiles of nuclear matrix proteins during the differentiation of human gastric mucous adenocarcinoma MGc80-3 cells. World J. Gastroenterol. 11, 4628–4633 (2005).

45. Murray, M. E., Mendez, M. G. & Janmey, P. A. Substrate stiffness regulates solubility of cellular vimentin. Mol. Biol. Cell 25, 87–94 (2014).

46. Soellner, P., Quinlan, R. A. & Franke, W. W. Identification of a distinct soluble subunit of an intermediate filament protein: tetrameric vimentin from living cells. Proc. Natl. Acad. Sci. U. S. A. 82, 7929–7933 (1985).

47. Premchandar, A. et al. Structural Dynamics of the Vimentin Coiled-coil Contact Regions Involved in Filament Assembly as Revealed by Hydrogen-Deuterium Exchange. J. Biol. Chem. 291, 24931–24950 (2016).

48. Mücke, N. et al. Molecular and biophysical characterization of assembly-starter units of human vimentin. J. Mol. Biol. 340, 97–114 (2004).

49. Keeling, M. C., Flores, L. R., Dodhy, A. H., Murray, E. R. & Gavara, N. Actomyosin and vimentin cytoskeletal networks regulate nuclear shape, mechanics and chromatin organization. Sci. Rep. 7, 5219 (2017).

50. Gesson, K. et al. A-type lamins bind both hetero- and euchromatin, the latter being regulated by lamina-associated polypeptide 2 alpha. Genome Res. 26, 462–473 (2016).

51. Pascual-Reguant, L. et al. Lamin B1 mapping reveals the existence of dynamic and functional euchromatin lamin B1 domains. Nat. Commun. 9, 3420 (2018).

52. Georgatos, S. D. & Blobel, G. Lamin B constitutes an intermediate filament attachment site at the nuclear envelope. J. Cell Biol. 105, 117–125 (1987).

53. Colucci-Guyon, E., Giménez Y Ribotta, M., Maurice, T., Babinet, C. & Privat, A. Cerebellar defect and impaired motor coordination in mice lacking vimentin. Glia 25, 33–43 (1999).

54. Boyne, L. J., Fischer, I. & Shea, T. B. Role of vimentin in early stages of neuritogenesis in cultured hippocampal neurons. Int. J. Dev. Neurosci. Off. J. Int. Soc. Dev. Neurosci. 14, 739–748 (1996).

55. Perlson, E. et al. Vimentin-Dependent Spatial Translocation of an Activated MAP Kinase in Injured Nerve. Neuron 45, 715–726 (2005).

56. Cavaluzzi, M. J. & Borer, P. N. Revised UV extinction coefficients for nucleoside-5’-monophosphates and unpaired DNA and RNA. Nucleic Acids Res. 32, e13–e13 (2004).

57. Abmayr, S. M., Yao, T., Parmely, T. & Workman, J. L. Current protocols in molecular biology/edited by Frederick M. Ausubel Al (2006).

58. Shevchenko, A., Wilm, M., Vorm, O. & Mann, M. Mass Spectrometric Sequencing of Proteins from Silver-Stained Polyacrylamide Gels. Anal. Chem. 68, 850–858 (1996).

